# Bioactivity-Driven Prediction of Antibacterial Synergy Using Machine Learning Models

**DOI:** 10.64898/2025.12.14.694088

**Authors:** Hannie Yousefabadi, Mahya Mehrmohamadi

**Affiliations:** Department of Molecular Biotechnology, School of Biotechnology, College of Science, University of Tehran, Street, Postcode, Tehran, Iran

**Keywords:** Antibacterial drug combinations, Synergy prediction, Antibacterial resistance, Nested cross-validation, Bioactivity signatures, Chemical Checker

## Abstract

**Motivation:** Predicting antibacterial drug synergy remains difficult due to strain variability and the limited scale of experimentally tested combinations. Existing machine-learning approaches often rely on permissive cross-validation schemes that allow drug pairs to appear across folds, inflating performance. A rigorous evaluation framework and scalable feature representation are needed for robust generalization.

**Results:** We assembled a curated dataset of 3,160 drug–pair–strain interactions covering 97 compounds and 10 bacterial strains. We then developed HALO (Held-out Antibiotic interaction Learning from latent bioactivity Observations), a synergy-prediction framework in which each drug pair is encoded using multi-level Chemical Checker (CC) similarity features spanning chemical, target, network, cellular, and clinical bioactivity domains. Under strictly nested, pair-heldout cross-validation (CV1), HALO achieved stable generalization to unseen combinations (accuracy ≈ 0.75; ROC–AUC = 0.82). Performance depended strongly on evaluation stringency: models performed well under random splits but degraded when required to generalize to unseen drug pairs and strain contexts. Despite these constraints, HALO generalized to an independent set of Loewe-*α* measurements, achieving ROC–AUC = 0.85 for distinguishing synergy from antagonism. These results demonstrate that multi-level bioactivity signatures provide a scalable, interpretable basis for predicting antibacterial synergy and reveal the performance limits of current models under rigorous evaluation.

**Availability and Implementation:** Code, data-processing scripts, and trained models will be available at GitHub repo.

**Supplementary information:** Supplementary figures and additional evaluation details are available online.

## 1. Introduction

Antimicrobial resistance (AMR) continues to rise globally and now poses a major public-health threat, with resistance spreading faster than new antibiotics are being developed [1, 2, 3]. Because the clinical pipeline remains dominated by derivatives of existing classes, combination therapy has become an important strategy for enhancing efficacy and slowing resistance emergence [4]. Yet antibiotic interactions are highly variable: synergy can improve killing and suppress resistance, whereas antagonism can reduce efficacy or promote failure [5, 6, 7]. Measuring these interactions at scale remains challenging—synergy assays are labor-intensive, context-dependent, and often vary across strains and conditions [8]. These limitations have motivated computational approaches for prioritizing combinations. While machine-learning models have shown strong performance in oncology [9, 10], antibacterial applications are hindered by sparse, heterogeneous datasets and many existing methods struggle to generalize to unseen drug pairs [11, 12]. The Chemical Checker (CC) offers a richer alternative to structure-only features by integrating chemical, biological, and phenotypic drug signatures into standardized embedding spaces [13]. Models that incorporate these bioactivity features consistently outperform purely structural descriptors [14], and CC-based frameworks—including CCSynergy, Bio-Mol, and recent repurposing studies—demonstrate that multi-level bioactivity embeddings can capture complex drug behavior relevant to combination modeling [15, 16, 17, 18]. However, CC-style embeddings have not been systematically applied to antibacterial interactions. Existing antibacterial models integrate chemical and biological information [19], but rely on small, heterogeneous datasets and rarely incorporate strain-level variation despite strong evidence that interaction outcomes differ across species, physiology, and strains [8, 12]. As a result, generalization to unseen drug pairs or unseen strains remains largely untested. To address these gaps, we curated 3,160 antibacterial pair–strain interactions from high-throughput screens and targeted literature sources and represented each drug using CC-based bioactivity signatures. We evaluated models under strict pair-held-out cross-validation with nested hyperparameter tuning. Our primary model, HALO-CV1, excludes strain embeddings yet generalizes reliably to unseen drug pairs. Comparisons with strain-aware variants (HALO-CV1s, HALO-CV2s) and a lenient baseline (HALO-ST) show that evaluation rigor strongly affects reported accuracy, and that CC-derived bioactivity features provide the dominant signal. Feature-importance analyses reveal biologically meaningful sublevels that shape predictions. Overall, our results demonstrate the feasibility of scalable, bioactivity-driven synergy prediction and establish a foundation for integrating richer strain-level or mechanistic representations in future work.

## 2. Results

### 2.1 Dataset construction and curation

We integrated interaction measurements from Brochado et al. [20], Cacace et al. [21], and manually curated literature-reported pairs from the ACDB [22] (Figure 1A). After standardizing drug identities and strains and restricting the compound space to approved antibacterials, Bliss *ε* values were harmonized across studies and consolidated using a reproducible procedure for resolving replicate inconsistencies. This yielded 3,160 high-quality pair–strain interactions spanning diverse drug classes and both Gram-positive and Gram-negative species (Figure 1B). Each compound was then encoded using multi-level Chemical Checker bioactivity signatures to generate pairwise similarity features (Figure 1C), providing a consistent representation of chemical, biological, and phenotypic drug properties. Together, the curated dataset and CC-derived features establish a standardized foundation for evaluating antibacterial synergy prediction models.

**Fig. 1.**
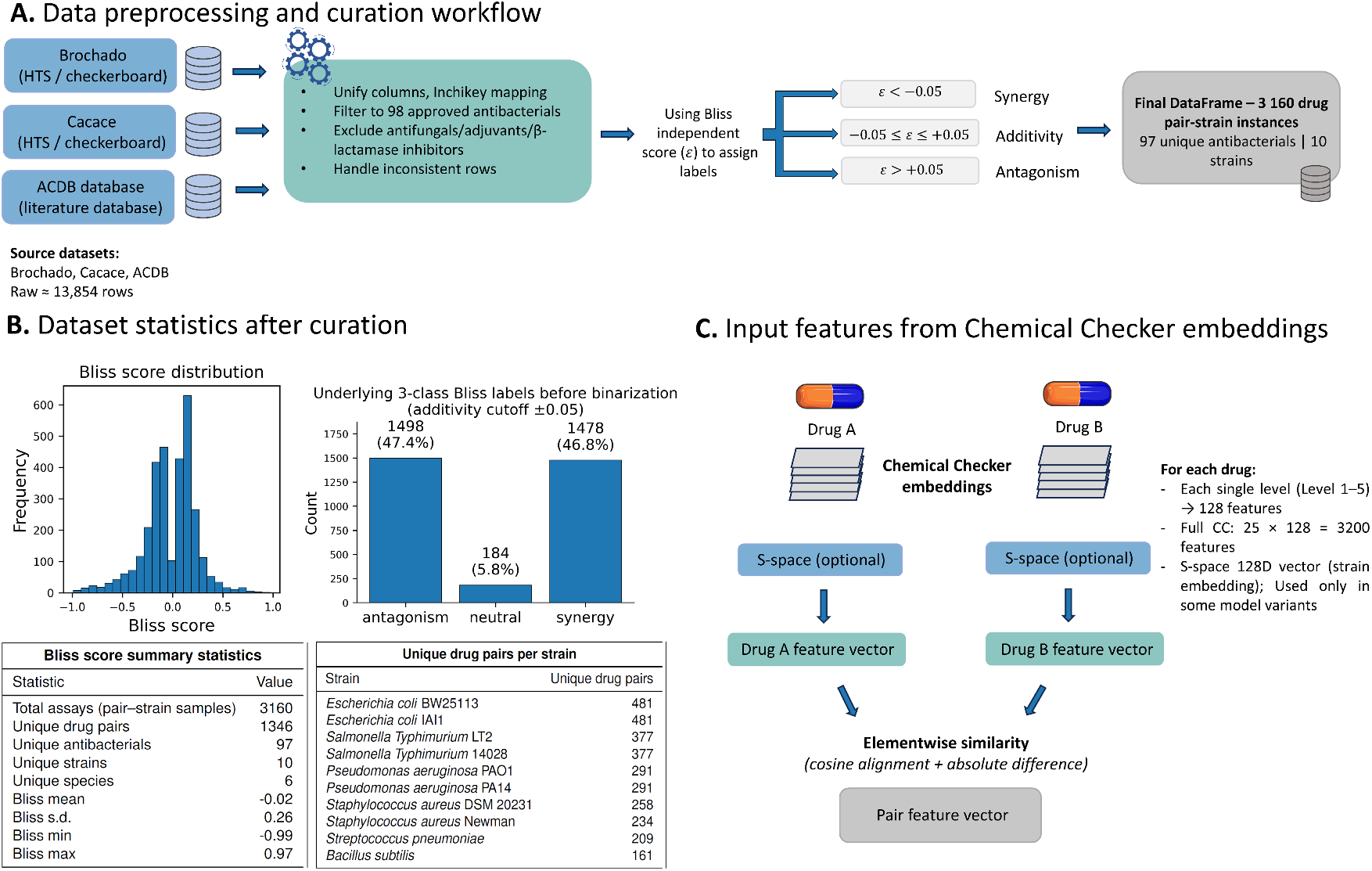
Data curation, dataset characteristics, and feature construction. (A) Workflow for preprocessing and integrating interaction data from sources. Interaction data were standardized, mapped to InChIKeys, filtered to 97 antibacterials, and cleaned for duplicates/inconsistencies. Bliss labels were assigned using *ε* < -0.05 (synergy) and *ε* > +0.05 (antagonism), yielding 3,160 pair–strain measurements across 10 strains. (B) Bliss score distribution, underlying 3-class labels before binarization, and summary statistics for strains, antibacterials, and score ranges. (C) Pairwise features derived from Chemical Checker embeddings: each drug’s 25×128-dimensional signature is converted into elementwise similarity features; an optional 128-dimensional strain embedding (S-Space) is included only in strain-aware models.

### 2.2 Performance of the HALO-CV1 model

We assessed HALO-CV1 using a five-fold pair-held-out scheme (CV1), reserving all interactions from 20% of drug pairs for testing and using nested three-fold group CV for hyperparameter selection (also grouped by drug pair to prevent data leakage). On the held-out set (N ≈ 602), HALO-CV1 achieved 0.75 accuracy, F1 scores in the 0.75–0.82 range, and a ROC–AUC of 0.82 (Figure 2A–D). The confusion matrix showed balanced performance across synergy and antagonism, with comparable false-positive and false-negative rates (Figure 2B). Performance was consistent across folds, with accuracy ≈0.72–0.77 and ROC–AUC ≈0.80–0.83 (Figure 2E), indicating reproducible generalization across distinct held-out drug-pair subsets. Overall, HALO-CV1—a simple, strain-agnostic model—generalizes reliably to unseen antibacterial combinations under strict pair-held-out evaluation.

**Fig. 2.**
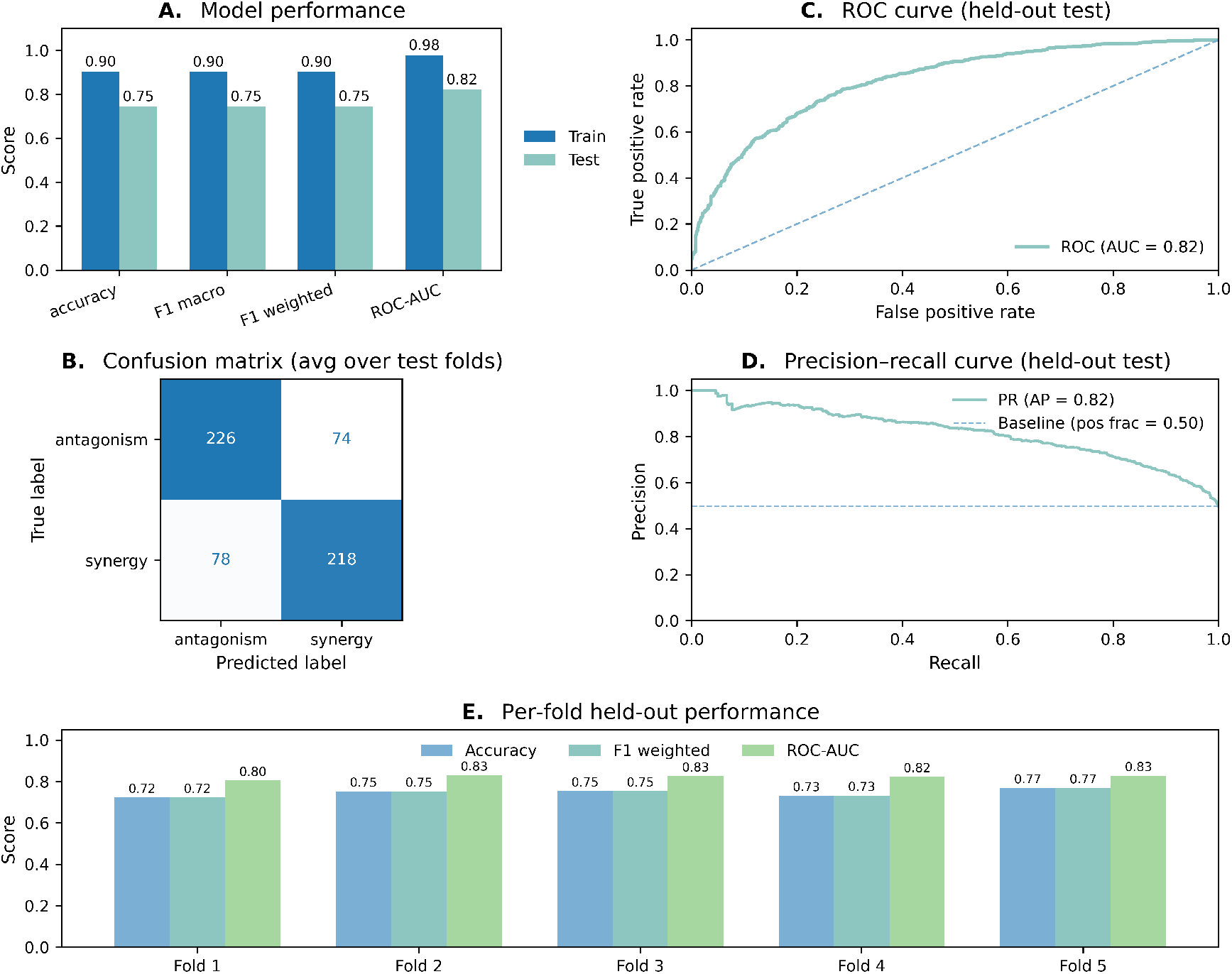
HALO-CV1 performance under pair-held-out cross-validation. (A) Train and outer-test performance (accuracy, F1-macro, F1-weighted, ROC-AUC) for CV1, where all drug pairs in the test fold are unseen. Test metrics range 0.75–0.82. (B) Average confusion matrix across outer folds (balanced classes). (C) ROC curve for held-out predictions (AUC = 0.82). The dotted line indicates random-chance performance. (D) Precision–recall curve (AP = 0.82; baseline = 0.50). (E) Per-fold outer-test accuracy, F1-weighted, and ROC-AUC, showing stable performance across all five CV1 folds (≈0.72–0.83).

### 2.3 Feature-level insights from Chemical Checker embeddings

We used LightGBM gain scores to identify which features contribute most to HALO-CV1’s predictions (Figure 3A–D). Individual elementwise CC similarities showed broadly distributed importance, with several chemical-genetic and phenotypic dimensions ranking highest (Figure 3A). When aggregated, CC sublevels tied to chemical-genetics, mechanism of action, small-molecule roles, signaling pathways, and side-effect profiles carried the most cumulative gain (Figure 3B). At the five top-level CC domains, contributions were nearly uniform ( 17–21% each; Figure 3C), indicating that synergy prediction draws from a broad, multi-scale bioactivity representation rather than any single biological layer, for example, structural information. Adding strain-space embeddings had little effect: in HALO-CV1s, CC features accounted for 97.2% of total gain, whereas strain-space contributed only 2.8% (Figure 3D). Overall, synergy prediction is supported by diverse CC bioactivity modalities, and CC signatures—not strain embeddings— and that the HALO-CV1 model’s interpretability is driven by a consistent pattern of biologically meaningful CC sublevels.

**Fig. 3.**
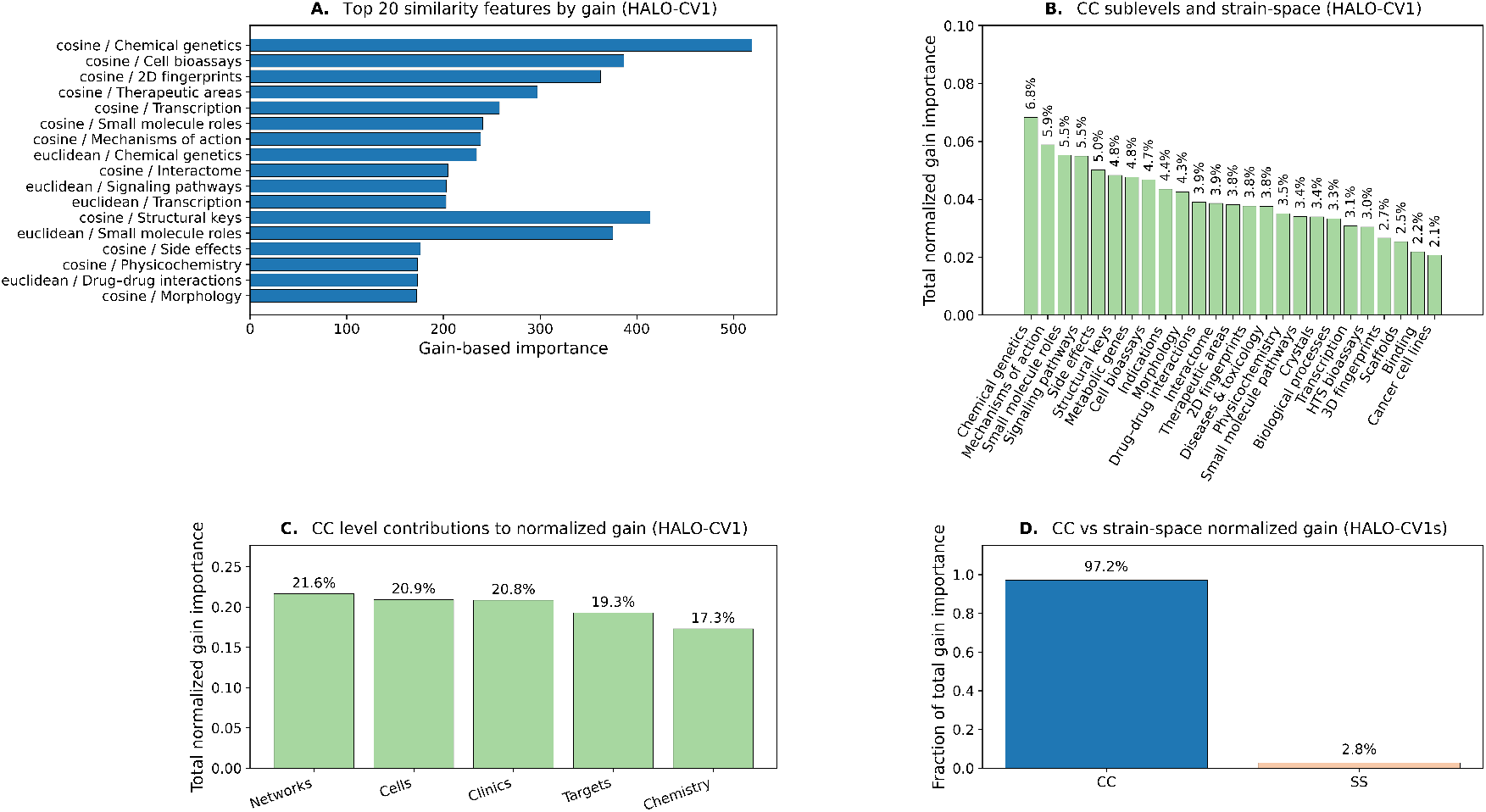
Feature contributions in HALO models. (A) Top 20 gain-ranked elementwise similarity features (HALO-CV1). (B) Normalized gain aggregated by Chemical Checker (CC) sublevels. (C) Total gain contributions from the five CC domains (Networks, Cells, Clinics, Targets, Chemistry). (D) Gain comparison in the strain-aware model (HALO-CV1s): CC features account for 97.2% of total gain, S-space for 2.8%.

### 2.4 External validation using the INDIGO antibacterial interaction dataset

We tested HALO-CV1 on an external dataset from Chandrasekaran et al. (INDIGO) [23], which reports Loewe *α* interaction scores for diverse antibiotic pairs in *E. coli* MG1655. Panel 4A summarizes the preprocessing pipeline used to map INDIGO compounds to InChIKeys, filter to approved antibacterials, attach CC similarity features, and assign synergy/antagonism labels from Loewe *α* scores (*α* < –0.5 synergy; *α* > 1 antagonism). After this curation, the resulting set showed strong class imbalance ( 19% synergy). Despite being trained only on Bliss-based interactions from different strains and conditions, HALO-CV1 achieved a ROC–AUC of 0.85 (Figure 4B). Average precision reached 0.56, well above the 0.19 baseline (Figure 4C), and the confusion matrix showed high specificity (47/57 antagonisms) with moderate synergy recall (9/13; Figure 4D). Predicted synergy probabilities also tracked monotonically with experimental *α* scores (Figure 4D). These results indicate that HALO-CV1 learns a transferable mapping from CC-derived similarity to antibacterial interaction outcomes, generalizing across assay types, strains, and data sources.

**Fig. 4.**
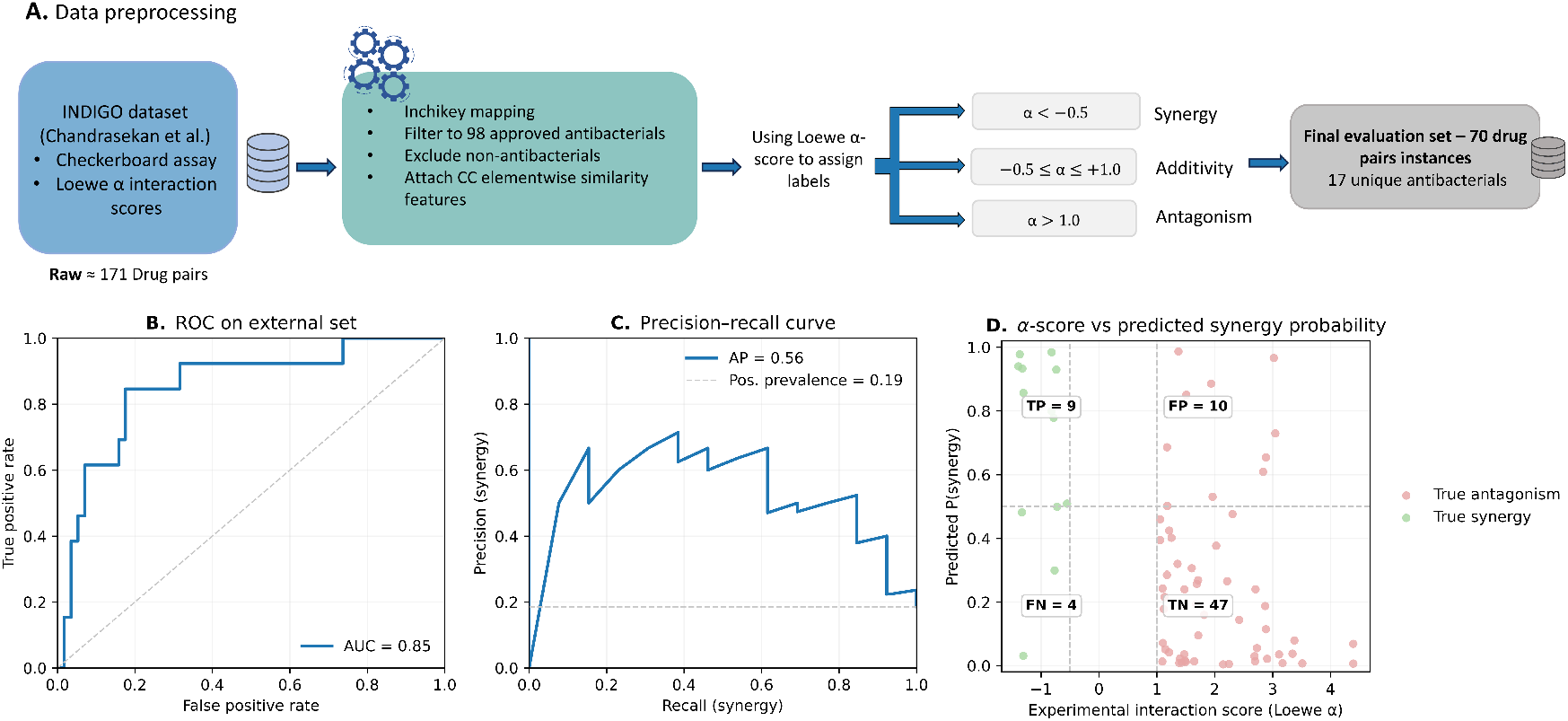
External validation on the INDIGO dataset. (A) External data preprocessing. The INDIGO dataset (≈171 pairs) was mapped to InChIKeys, filtered to approved antibacterials, non-antibacterials removed, and CC similarity features added. Loewe *α* thresholds (*α* < -0.5 synergy; *α* > 1 antagonism) defined labels. After filtering and excluding additive cases, 70 pairs (17 antibacterials) remained for external validation. (B) ROC curve for HALO-CV1 on 70 INDIGO pairs (AUC = 0.85). (C) Precision–recall curve (AP = 0.56; synergy prevalence = 0.19). Confusion outcomes: 47 TN, 9 TP, 10 FP, 4 FN. (D) Experimental Loewe *α* scores vs. predicted P(synergy), showing high probabilities for *α* < -0.5 (synergy) and low probabilities for *α* > 1 (antagonism).

### 2.5 Comparison of evaluation schemes and model variants

We evaluated all models using nested cross-validation with a 5-fold outer split (CV1: pair-held-out; CV2: pair+strain-held-out) and a 3-fold inner loop grouped by drug pair for hyperparameter selection (Supplementary Figure 1C). To contextualize HALO-CV1, we compared performance across validation schemes and model variants (Supplementary Figure 1). Under standard stratified CV (HALO-ST), where drug pairs can appear in multiple folds, performance was substantially inflated (accuracy/F1-macro ≈0.78; ROC–AUC ≈0.86). Enforcing pair-level separation (HALO-CV1s) reduced accuracy and F1 to ≈0.73 and ROC–AUC to ≈0.73–0.82, reflecting a more realistic generalization estimate. The strictest setting, holding out both drug pairs and strains (HALO-CV2s), yielded the lowest performance (≈0.53–0.58), consistent with the increased difficulty of predicting across strain shifts. Supplementary Figure 1B compares the two CV1 variants. HALO-CV1s (with strain-space embedding) performed equivalently to HALO-CV1 (CC-only), indicating that CC-derived pairwise similarities capture nearly all predictive signal under pair-held-out evaluation. Supplementary Figure 1C summarizes the nested CV workflow: the outer split defines the held-out fold, the inner loop selects hyperparameters, and the final model is refit on the full outer-training set before a single evaluation on the untouched outer-test fold. HALO-ST lacks this safeguard and therefore overestimates performance. Overall, stricter evaluation substantially lowers reported accuracy, HALO-CV1 matches the strain-aware variant under CV1, and CV1 provides a stable and realistic regime for assessing antibacterial synergy prediction.

### 2.6 Novel Drug-Pair Predictions and CC Similarity Patterns

Using the final HALO-CV1 model trained on all labeled interactions, we generated predictions for all drug pairs absent from the training set. Synergistic pairs consistently showed higher bioactivity concordance across CC levels (mean CC cosine ≈0.57; CC Euclidean ≈0.60), whereas antagonistic pairs exhibited a distinctly lower-similarity range (mean CC cosine ≈0.48; CC Euclidean ≈0.52) based on the top 20 highest-confidence predictions. Full ranked lists are provided in Supplementary Tables S2–S3 and top candidates in Table 1.

**Table 1.** Top 10 predicted synergistic drug combinations according to HALO-CV1.

| Top 10 Predicted Synergistic Pairs (HALO-CV1) |  |  |  |  |  |
| --- | --- | --- | --- | --- | --- |
| Drug Pair | P(syn) | P(ant) | Drug Classes (A-B) | CC Cosine Similarity | CC Euclidean Similarity |
| Bacitracin + Vancomycin | 0.998 | 0.002 | Polypeptide antibacterial – Glycopeptide | 0.599 | 0.724 |
| Bacitracin + Oritavancin | 0.997 | 0.003 | Polypeptide antibacterial – Glycopeptide (lipoglycopeptide) | 0.653 | 0.711 |
| Mecillinam + Cefuroxime | 0.995 | 0.005 | Penicillin (amidino) – Cephalosporin (2nd gen) | 0.627 | 0.693 |
| Bacitracin + Minocycline | 0.994 | 0.006 | Polypeptide antibacterial – Tetracycline-class antibiotic | 0.543 | 0.620 |
| Cefuroxime + Penicillin G | 0.993 | 0.007 | Cephalosporin (2nd gen) – Natural penicillin | 0.655 | 0.660 |
| Bacitracin + Thymol | 0.993 | 0.007 | Polypeptide antibacterial – Monoterpene phenol | 0.456 | 0.487 |
| Cefuroxime + Amikacin | 0.993 | 0.007 | Cephalosporin (2nd gen) – Aminoglycoside | 0.545 | 0.596 |
| Bacitracin + Procaine | 0.992 | 0.008 | Polypeptide antibacterial – Local anesthetic (ester) | 0.463 | 0.561 |
| Bacitracin + Daptomycin | 0.992 | 0.008 | Polypeptide antibacterial – Lipopeptide antibiotic | 0.762 | 0.759 |
| Telithromycin + Tedizolid | 0.991 | 0.009 | Ketolide – Oxazolidinone | 0.477 | 0.533 |

## 3. Methods

### 3.1 Data sources and antibacterial curation

Interaction data were compiled from three sources: high-throughput checkerboard assays from Brochado et al. [20] and Cacace et al. [21], and manually curated pairwise interaction entries from the ACDB database [22]. To define a consistent antibacterial compound space, we constructed a unified list of antibacterial compounds by integrating FDA-approved agents with research and literature-reported antibacterial compounds. Non-antibacterial agents, combination products, and *β*-lactamase inhibitors were excluded. All compound identities were standardized by mapping names and stereochemical variants to canonical InChIKeys. The dataset spans 10 commonly used Gram-positive and Gram-negative strains, including *E. coli* BW25113/ IAI1, *S. Typhimurium* 14028/ LT2, *P. aeruginosa* PA14/ PAO1, *S. aureus* DSM20231/ Newman, *S. pneumoniae*, and *B. subtilis*, enabling evaluation across diverse physiological contexts.

### 3.2 Interaction score harmonization and dataset integration

All datasets reported Bliss *ε* values but differed in checkerboard resolution and aggregation. After removing exact duplicates, replicated measurements for the same drug–strain pair were merged using a reproducible rule: exact replicates were collapsed; near-consistent values (|Δ*ε*| ≤ 0.05) were averaged; and conflicting sets were curated by discarding the farthest-from-median value before averaging. Rows with missing essential identifiers were excluded. Each (Drug A, Drug B, Strain) combination was represented by one harmonized *ε* score, yielding 3,160 interactions across 97 antibacterials and 10 strains. To obtain reliable binary labels, we excluded interaction values close to the additive boundary. Although Bliss *ε* values between -0.1 and +0.1 are commonly considered “neutral,” the distribution of our curated dataset showed high variability and reduced separability near this range (Supplementary Figure 2). To increase label purity, we applied a margin-based threshold and removed samples with |*ε*| ≤ 0.05. Interactions with *ε* < -0.05 were labeled as synergy and *ε* > +0.05 as antagonism. This margin-based relabeling is consistent with standard practices for handling noisy or continuous interaction metrics in machine learning.

### 3.3 Feature construction

For each drug, we retrieved sig2 Chemical Checker (CC) embeddings (25 × 128-dimensional subspaces). Pairwise drug features were computed using elementwise cosine similarity and Euclidean-derived similarity (1 - normalized L2 distance). Across 25 CC spaces, this produced 6,400 raw features per drug pair. Because CC embeddings are bounded and internally normalized, no additional feature scaling was applied. A strain-space embedding derived from strain–drug response profiles was also generated using the Chemical Checker protocol [24]. This additional feature set was used only in the strain-aware HALO-CV1s and HALO-CV2s models. A reduced feature subset derived from LightGBM-based feature selection served as the input representation for all models. This feature selection step yielded 1,578 CC-based pairwise similarity features, and this fixed feature set was used in HALO-CV1 and HALO-ST. Although the same LightGBM-based feature-selection framework was also applied when constructing the strain-aware HALO-CV1s and HALO-CV2s variants, the exact set and number of selected features differ across those models, reflecting differences in input dimensionality and grouping schemes. For all binary classification tasks, interaction labels were defined as synergy (*ε* < –0.05) and antagonism (*ε* > 0.05), with neutral interactions excluded prior to modeling.

### 3.4 Model training and cross-validation

All models were implemented using LightGBM for binary classification, with “synergy” treated as the positive class. Training followed a nested cross-validation design to ensure strict separation between model selection and evaluation. The outer split defined the held-out test set and was constructed using one of two grouping schemes. In the CV1 (pair-held-out) scheme, five outer folds were generated with a StratifiedGroupKFold splitter grouping by drug pair, such that all measurements for 20% of drug pairs were withheld together in each fold. The CV2 (pair + strain–held-out) scheme used five independent brute-force outer splits, each selecting a subset of strains whose induced drug-pair set yielded a test size within predefined bands (16–28% of all rows). For each split, both strains and drug-pair combinations present in the test subset were completely excluded from training, and any cross-edge rows violating this disjointness were removed. Hyperparameter tuning occurred strictly inside the training portion of each outer split, using a three-fold StratifiedGroupKFold grouped by drug pair. Thirty-two LightGBM configurations were sampled from a predefined search space, trained with early stopping for stable convergence, and ranked by mean inner-CV accuracy. The best configuration was then retrained on the full outer-training set and evaluated once on the untouched outer-test set. For comparison, a lenient standard stratified cross-validation baseline (HALO-ST) was included; because it does not enforce pair-level grouping or nested evaluation, it provides optimistically biased estimates.

### 3.5 Evaluation metrics and feature importance analysis

Performance was assessed using accuracy, F1-macro/weighted, ROC–AUC, and PR–AUC. Confusion matrices used synergy as the positive class. Because the test sets were approximately balanced between synergy and antagonism, random-chance baselines for PR–AUC were 0.50. Feature importance in the main HALO-CV1 model was quantified using LightGBM gain-based importance, which reflects each feature’s contribution to the reduction of loss across decision tree splits. Importances were normalized and aggregated across CC sublevels and high-level CC domains. For HALO-CV1s, the contribution of strain-derived features was compared with CC-derived features.

### 3.6 Prediction of novel and external pairs

The final HALO-CV1 model was retrained on the complete labeled dataset (1,578 selected CC features, 3160 drug-pair-strain combinations) and used to score all possible unlabeled drug–drug pairs. High-confidence predicted synergistic and antagonistic pairs were reported. External validation used the INDIGO dataset of Chandrasekaran et al. [23], which reports quantitative Loewe *α* interaction scores for 171 *E. coli* pairs. After mapping compounds, label binarization, excluding non-antibacterials, and retaining only pairs with CC coverage, 70 pairs remained. Labels were binarized (*α* < –0.5 = synergy; *α* > 1 = antagonism). CC-based elementwise similarity features for this external set were generated with the same FeatureMapper pipeline and restricted to the exact same 1,578 features used by HALO-CV1. The final HALO-CV1 model (trained on all internal data with fixed hyperparameters) was then applied to these 70 combinations, and external performance was evaluated identically to internal tests.

### 3.7 Software and reproducibility

All analyses were performed in Python 3.12.3 on a Linux-based high-performance server environment. Model training used LightGBM 4.6.0 and scikit-learn 1.7.2, pandas 2.3.3, numpy 2.3.5. and matplotlib 3.10.3. CC signatures were retrieved via the Chemical Checker API. Random seeds were fixed for all steps. Custom scripts and data are publicly available at (GitHub repository link to be added). Additional methodological details, including alternative feature encodings, exploratory model variants, and neural-network baselines, are provided in the Supplementary Methods.

## 4. Discussion

### 4.1. Summary of main findings

In this work we assembled a curated resource of 3,160 antibacterial drug–pair–strain interactions and used Chemical Checker–derived bioactivity similarities to train supervised models for synergy prediction. Under a strict pair-held-out evaluation (CV1), our main model, HALO-CV1, achieved stable performance on unseen drug combinations (accuracy ≈0.75, ROC–AUC ≈0.82) while maintaining balanced precision and recall for both synergistic and antagonistic classes. These results show that multi-level bioactivity signatures, without any explicit strain embedding, are sufficient to capture a substantial fraction of the variability in antibacterial interaction outcomes.

### 4.2. Relation to previous experimental and computational work

High-throughput screening studies have shown that antibacterial drug interactions are common yet strongly context-dependent, with most combinations clustering near additivity and true synergy or antagonism occurring infrequently across species and strains [20, 21, 25]. This variability, together with limited strain coverage and noisy measurements, has constrained the development and evaluation of predictive models for antibacterial synergy. Existing computational approaches can be broadly grouped into three categories. Network- and graph-based models leverage drug–target or protein–interaction networks, but are typically trained on small datasets and evaluated under random or weakly stratified cross-validation, limiting assessment of generalization to unseen drug pairs. Metabolism- and flux-based methods integrate genome-scale metabolic models with machine learning to predict interaction outcomes, offering mechanistic insight but focusing on condition-specific settings rather than pair-level extrapolation. Structure-based chemoinformatic models, such as CoSynE, use molecular fingerprints and have shown enrichment of validated synergies in prospective tests, yet similarly rely on evaluation schemes that allow substantial overlap between training and test drug pairs [26]. Among antibacterial-specific frameworks, INDIGO combines chemogenomic profiles with random forests to predict interactions in *E. coli*, achieving moderate accuracy and partial transferability across species via orthology mapping [23]. Lv et al. graph-learning model outperforms INDIGO and CoSynE with precision ≈ 0.875 and accuracy of 0.90 [27]. M2D2 machine learning framework extended prediction to two and three drug combination predictions and achieved accuracy ≈ 0.82 on evaluation set [28]. However, across these methods, evaluation is generally performed under random or partially stratified cross-validation within a single strain, where many drug pairs—or close analogs—appear in both training and test sets. As a result, reported accuracies primarily reflect interpolation rather than true generalization to unseen combinations. Our work differs in two key respects. First, we focus on curated, multi-strain antibacterial interaction prediction, enabling direct assessment of model generalization across biological contexts rather than within a single strain. Second, we evaluate performance under strictly nested cross-validation, explicitly separating hyperparameter optimization from final evaluation. We further impose pair-level and pair-plus-strain-level holdout schemes, allowing us to quantify how predictive performance degrades as evaluation stringency increases. Under pair-level generalization (CV1), HALO-CV1 achieves accuracy and F1-macro of approximately 0.75 and ROC–AUC of 0.82, despite substantially higher training performance. This gap highlights the extent to which lenient validation schemes overestimate model performance. Building on prior Chemical Checker–based frameworks [15, 16, 17, 18], we extend multi-level bioactivity similarity to antibacterial interactions. Our contribution lies not in new representations but in a rigorous evaluation framework that distinguishes pair memorization from transferable similarity, providing a more conservative and biologically meaningful benchmark for antibacterial synergy prediction.

### 4.3. External validation and biological interpretation of predicted novel pairs

To assess generalization beyond our curated dataset, we evaluated HALO-CV1 on the INDIGO interaction measurements, which differ in assay type, interaction scoring, and drug–pair composition. Despite these shifts, HALO-CV1 achieved a ROC–AUC of 0.85 (Fig4B) and produced synergy scores that varied monotonically with experimental Loewe *α* values (Fig4D), indicating robust ranking of interaction strength across assay modalities [23]. High-confidence synergistic pairs occupy regions of elevated Chemical Checker similarity, while antagonistic pairs cluster at intermediate similarity, consistent with prior mechanistic models of antibacterial interactions; it mirrors observations from Brochado et al. and Cacace et al., where synergies tended to arise between drugs acting on the same biological process and antagonisms between drugs targeting different processes [20, 21]. The intermediate-similarity regime associated with antagonism is likewise consistent with Lehár et al.’s topology model, in which partially overlapping pathways buffer each other’s effects [29]. We next highlight representative high-confidence predictions to illustrate how HALO-CV1 aligns with known mechanisms or suggests biologically plausible new combinations. Several high-confidence predictions align with known or class-level mechanisms. For example, Bacitracin–Vancomycin and Bacitracin–Oritavancin combine agents that disrupt cell-wall synthesis and membrane integrity respectively, providing a coherent rationale for synergy despite the absence of direct studies for some pairs [30]. The predicted Mecillinam–Cefuroxime interaction is also biologically plausible: Mecillinam synergizes with related cephalosporins such as Cephradine against Gram-negative species [31], and Mecillinam-resistant CTX-M-15 mutants exhibit collateral sensitivity to Cefuroxime [32], supporting its plausibility. Other predictions extend established class-level precedents. While Cefuroxime has not been evaluated with Penicillin G, strong synergy has been reported between Cefuroxime and the penicillin-class agent Ticarcillin across multiple Gram-negative pathogens [33]. Similarly, the Cefuroxime–Amikacin prediction is consistent with extensive evidence for *β*-lactam–aminoglycoside synergy, including 77.5% synergy for Ceftriaxone–Amikacin in *P. aeruginosa* and confirmed in vivo activity [34, 35]. HALO-CV1 also identifies less intuitive but biologically coherent hypotheses. The Bacitracin–Minocycline prediction (P(syn)=0.994) would traditionally be expected to show antagonism based on bactericidal–bacteriostatic reasoning, yet is supported by strong phenotypic similarity encoded in the Chemical Checker representation. Additional untested predictions, including Bacitracin–Thymol, Bacitracin– Daptomycin, and Telithromycin–Tedizolid, are consistent with envelope-targeting mechanisms or complementary ribosomal inhibition, and therefore represent fully model-driven hypotheses for experimental follow-up [36, 37].

### 4.4. Scope of the model: pair generalization vs. strain generalization

When required to generalize simultaneously to unseen drug pairs and unseen strains, performance drops toward chance in the strain-aware HALO-CV2s setting. This reflects data sparsity rather than model failure: only ten strains are available, most drug pairs are observed in at most one strain, and pair–strain overlap is minimal. In this regime, explicit strain embeddings add little beyond Chemical Checker–derived bioactivity similarity, which already captures much of the transferable mechanistic signal. Accordingly, HALO-CV1 is best interpreted as a tool for prioritizing new drug pairs in well-characterized strain panels, matching typical experimental screening settings.

### 4.5. Data limitations, challenges, and future directions

Our results reveal factors that constrain predictive performance. High-quality interaction data remain limited: most Bliss measurements lie near additivity, are noisy around *ε* ≈ 0, and cover only a small set of strains, reducing the model’s ability to learn strain-specific patterns. Dose dependence poses an additional challenge, as our predictor relies on dose-aggregated *ε* values from high-throughput screens and heterogeneous literature reports, preventing it from capturing concentration-specific interaction shifts. Finally, sparse overlap of drug pairs across strains limits the feasibility of strain-aware modeling, as seen in the poor generalization of HALO-CV2s. Progress will require broader and more standardized multi-strain datasets, as well as richer strain-level features such as genomic or metabolic profiles. Hybrid models that integrate CC bioactivity embeddings with mechanistic representations may also improve accuracy and interpretability. Overall, CC-based similarities provide a solid foundation under current data constraints, but larger, better-resolved datasets are essential for advancing truly strain-aware prediction.

## 5. Conclusion

In this study, we curated 3,160 antibacterial drug–pair–strain interactions and developed a Chemical Checker–based framework for predicting antibiotic synergy. Under strict pair-held-out evaluation with nested cross-validation, our primary model, HALO-CV1, achieved stable and balanced performance on unseen drug combinations, demonstrating that multi-level bioactivity signatures alone provide a strong and interpretable basis for antibacterial interaction prediction. Across evaluation schemes, we found that cross-validation strictness has a major impact on apparent performance. The strain-aware HALO-CV2s model performs poorly when required to generalize simultaneously to unseen drug pairs and unseen strains, reflecting current data limitations rather than a failure of the modeling approach. With sparse strain coverage and limited replication of pair–strain combinations, strain embeddings add little beyond the bioactivity similarity captured by CC features, which currently dominate predictive signal. These findings clarify the practical scope of our approach. HALO-CV1 is most suitable for prioritizing new drug combinations in well-characterized laboratory or clinical strains—an evaluation setting that aligns with many experimental screening workflows. At the same time, our analysis highlights the need for broader multi-strain datasets, deeper strain-level characterization, and integration of richer omics or mechanistic features to support future cross-strain generalization. The strong performance of HALO-CV1 across independent experimental platforms further underscores the robustness of the underlying CC representation, while its evaluation under rigorously nested pair- and pair–strain-held-out schemes provides a more realistic benchmark than prior work. Overall, this study establishes a reproducible baseline for antibacterial synergy prediction and a framework for benchmarking future models under data-realistic evaluation conditions. As larger and more diverse datasets emerge, this foundation will support increasingly accurate and mechanistically grounded computational discovery of antibacterial drug combinations.

## Competing interests

No competing interest is declared.

## Author contributions statement

HY: data curation, methodology, formal analysis, original draft. MM: conceptualization, supervision, review and editing.

## Acknowledgement

We thank Milad Reyhani for his support and contributions to data acquisition for this work. Image(s) provided by Servier Medical Art (https://smart.servier.com), licensed under CC BY 4.0 (https://creativecommons.org/licenses/by/4.0/). AI Disclosure: Large language models (ChatGPT 5.1 and Opus 4.5) were used to help edit and polish portions of the manuscript text. The authors revised and approved all content and are responsible for the scientific accuracy of the work.

## Notes

### Competing Interest Statement

The authors have declared no competing interest.

## References

1. Md. Abdus Salam, Md. Yusuf Al-Amin, Moushumi Tabassoom Salam, Jogendra Singh Pawar, Naseem Akhter, Ali A. Rabaan, and Mohammed A. A. Alqumber. Antimicrobial Resistance: A Growing Serious Threat for Global Public Health. Healthcare, 11(13):1946, July 2023.

2. Antimicrobial resistance.

3. Daniela Melchiorri, Tamarie Rocke, Richard A Alm, Alexandra M Cameron, and Valeria Gigante. Addressing urgent priorities in antibiotic development: insights from WHO 2023 antibacterial clinical pipeline analyses. The Lancet. Microbe, 6(3):None, March 2025.

4. Bence Bognár, Réka Spohn, and Viktória Lázár. Drug combinations targeting antibiotic resistance. npj Antimicrobials and Resistance, 2:29, October 2024.

5. J. F. Acar. Antibiotic synergy and antagonism. The Medical Clinics of North America, 84(6):1391–1406, November 2000.

6. R. C. Moellering. Rationale for use of antimicrobial combinations. The American Journal of Medicine, 75(2A):4–8, August 1983.

7. G. M. Eliopoulos and C. T. Eliopoulos. Antibiotic combinations: should they be tested? Clinical Microbiology Reviews, 1(2):139–156, April 1988.

8. Antimicrobial interactions: mechanisms and implications for drug discovery and resistance evolution - ScienceDirect.

9. Machine learning approaches for drug combination therapies - PMC.

10. Harnessing machine learning to find synergistic combinations for FDA-approved cancer drugs | Scientific Reports.

11. A systematic evaluation of deep learning methods for the prediction of drug synergy in cancer | PLOS Computational Biology.

12. Jennifer M. Cantrell, Carolina H. Chung, and Sriram Chandrasekaran. Machine learning to design antimicrobial combination therapies: Promises and pitfalls. Drug Discovery Today, 27(6):1639–1651, June 2022.

13. Extending the small-molecule similarity principle to all levels of biology with the Chemical Checker | Nature Biotechnology.

14. Srijit Seal, Hongbin Yang, Maria-Anna Trapotsi, Satvik Singh, Jordi Carreras-Puigvert, Ola Spjuth, and Andreas Bender. Merging bioactivity predictions from cell morphology and chemical fingerprint models using similarity to training data. Journal of Cheminformatics, 15(1):56, June 2023.

15. CCSynergy: an integrative deep-learning framework enabling context-aware prediction of anti-cancer drug synergy | Briefings in Bioinformatics | Oxford Academic.

16. Hiu Fung Yip, Xiao Wei, Zeming Li, Qing Ren, DongSheng Cao, Lu Zhang, and Aiping Lu. Bio-Mol:Pretraining Multimodality Bioactivity Profile for Enhancing Small Molecule Property Prediction, November 2023. Pages: 2023.11.02.565401 Section: New Results.

17. Gong-Hua Li, Feifei Han, Rong-Hua Luo, Peng Li, Chia-Jung Chang, Weihong Xu, Xin-Yan Long, Jing-Fei Huang, Yong-Tang Zheng, Qing-Peng Kong, and Wenzhong Xiao. Predictive Systems Biology Modeling: Unraveling Host Metabolic Disruptions and Potential Drug Targets in Acute Viral Infections, August 2023. Pages: 2023.07.24.550423 Section: New Results.

18. Eva Viesi, Ugo Perricone, Patrick Aloy, and Rosalba Giugno. APBIO: bioactive profiling of air pollutants through inferred bioactivity signatures and prediction of novel target interactions. Journal of Cheminformatics, 17(1):13, January 2025.

19. Lei Chen, Bi-Qing Li, Ming-Yue Zheng, Jian Zhang, Kai-Yan Feng, and Yu-Dong Cai. Prediction of Effective Drug Combinations by Chemical Interaction, Protein Interaction and Target Enrichment of KEGG Pathways. BioMed Research International, 2013:723780, 2013.

20. Species-specific activity of antibacterial drug combinations | Nature.

21. Elisabetta Cacace, Vladislav Kim, Vallo Varik, Michael Knopp, Manuela Tietgen, Amber Brauer-Nikonow, Kemal Inecik, André Mateus, Alessio Milanese, Marita Torrissen Mårli, Karin Mitosch, Joel Selkrig, Ana Rita Brochado, Oscar P. Kuipers, Morten Kjos, Georg Zeller, Mikhail M. Savitski, Stephan Göttig, Wolfgang Huber, and Athanasios Typas. Systematic analysis of drug combinations against Gram-positive bacteria. Nature Microbiology, 8(11):2196–2212, November 2023. Publisher: Nature Publishing Group.

22. Frontiers | ACDB: An Antibiotic Combination DataBase.

23. Sriram Chandrasekaran, Melike Cokol-Cakmak, Nil Sahin, Kaan Yilancioglu, Hilal Kazan, James J. Collins, and Murat Cokol. Chemogenomics and orthology-based design of antibiotic combination therapies. Molecular Systems Biology, 12(5):872, May 2016.

24. Arnau Comajuncosa-Creus, Martino Bertoni, Miquel Duran-Frigola, Adrià Fernández-Torras, Oriol Guitart-Pla, Nils Kurzawa, Martina Locatelli, Yasmmin Martins, Elena Pareja-Lorente, Gema Rojas-Granado, Nicolas Soler, Eva Viesi, and Patrick Aloy. Integration of diverse bioactivity data into the Chemical Checker compound universe. Nature Protocols, 20(11):3270–3294, November 2025.

25. Within-species variability of antibiotic interactions in Gram-negative bacteria | mBio.

26. Daniel J. Mason, Ian Stott, Stephanie Ashenden, Zohar B. Weinstein, Idil Karakoc, Selin Meral, Nurdan Kuru, Andreas Bender, and Murat Cokol. Prediction of Antibiotic Interactions Using Descriptors Derived from Molecular Structure. Journal of Medicinal Chemistry, 60(9):3902– 3912, May 2017.

27. Ji Lv, Guixia Liu, Yuan Ju, Ying Sun, and Weiying Guo. Prediction of Synergistic Antibiotic Combinations by Graph Learning. Frontiers in Pharmacology, 13, March 2022. Publisher: Frontiers.

28. Margaret M. Reuter, Katherine L. Lev, Jon Albo, Harkirat Singh Arora, Nemo Liu, Shenghao Tan, Madeline R. Shay, Debmalya Sarkar, Aaron Robida, David H. Sherman, Rudy J. Richardson, Nate J. Cira, and Sriram Chandrasekaran. Ultra-high-throughput screening of antimicrobial combination therapies using a two-stage transparent machine learning model. bioRxiv: The Preprint Server for Biology, page 2024.11.25.625231, November 2024.

29. Joseph Lehár, Grant R. Zimmermann, Andrew S. Krueger, Raymond A. Molnar, Jebediah T. Ledell, Adrian M. Heilbut, Glenn F. Short, Leanne C. Giusti, Garry P. Nolan, Omar A. Magid, Margaret S. Lee, Alexis A. Borisy, Brent R. Stockwell, and Curtis T. Keith. Chemical combination effects predict connectivity in biological systems. Molecular Systems Biology, 3(1):MSB4100116, February 2007.

30. R. Hancock and P. C. Fitz-James. SOME DIFFERENCES IN THE ACTION OF PENICILLIN, BACITRACIN, AND VANCOMYCIN ON BACILLUS MEGATERIUM. Journal of Bacteriology, 87(5):1044–1050, May 1964.

31. B. Chattopadhyay and I. Hall. In vitro combination of mecillinam with cephradine or amoxycillin for organisms resistant to single agents. The Journal of Antimicrobial Chemotherapy, 5(5):549–553, September 1979.

32. Carola E. H. Rosenkilde, Christian Munck, Andreas Porse, Marius Linkevicius, Dan I. Andersson, and Morten O. A. Sommer. Collateral sensitivity constrains resistance evolution of the CTX-M-15 β-lactamase. Nature Communications, 10(1):618, February 2019. Publisher: Nature Publishing Group.

33. C. S. Goodwin, W. E. S. Harper, and J. Epis. Synergy between cefuroxime and ticarcillin against Providencia, and the activity of cefuroxime against Streptococcus faecalis and gram-negative bacteria. Pathology, 12(2):294–295, January 1980.

34. P. Angehrn. In vitro and in vivo synergy between ceftriaxone and aminoglycosides against Pseudomonas aeruginosa. European Journal of Clinical Microbiology, 2(5):489–495, October 1983.

35. A classic antibiotic reimagined: Rationally designed bacitracin variants exhibit potent activity against vancomycin-resistant pathogens | PNAS.

36. Cristina Gan, Elisa Langa, Antonio Valenzuela, Diego Ballestero, and M. Rosa Pino-Otín. Synergistic Activity of Thymol with Commercial Antibiotics against Critical and High WHO Priority Pathogenic Bacteria. Plants, 12(9):1868, May 2023.

37. Sarah C J Jorgensen, Evan J Zasowski, Trang D Trinh, Abdalhamid M Lagnf, Sahil Bhatia, Noor Sabagha, Jacinda C Abdul-Mutakabbir, Sara Alosaimy, Ryan P Mynatt, Susan L Davis, and Michael J Rybak. Daptomycin Plus β-Lactam Combination Therapy for Methicillin-resistant Staphylococcus aureus Bloodstream Infections: A Retrospective, Comparative Cohort Study. Clinical Infectious Diseases, 71(1):1–10, June 2020.

