## Supplementary Figures for "Bioactivity-Driven Prediction of Antibacterial Synergy Using Machine Learning Models"

### Supplementary Fig1:

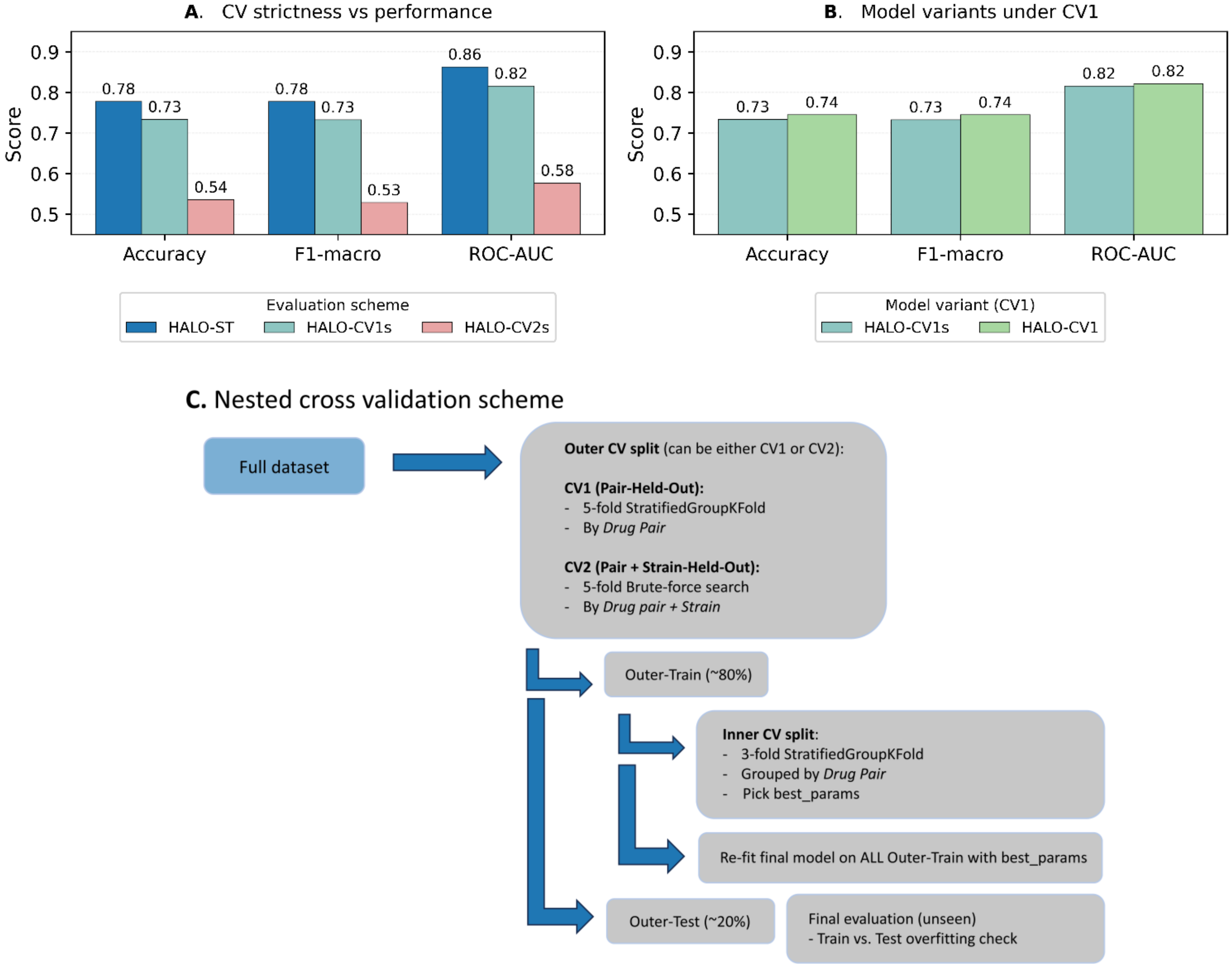


**Supplementary Figure 1. Comparison of validation schemes and model variants.**

***(A)*** *Performance of the same model under three evaluation schemes: standard stratified CV (HALO-ST), pair-held-out CV (HALO-CV1s), and pair+strain-held-out CV (HALO-CV2s).* ***(B)*** *Comparison of HALO-CV1 (CC features only) vs. HALO-CV1s (CC + strain-space) under the CV1 split.* ***(C)*** *Nested CV workflow: a 5-fold outer split (CV1 or CV2) defines held-out data; a 3-fold inner CV grouped by drug* *pair tunes hyperparameters; the final model is refit on the outer-train fold and evaluated once on the outer-test fold.*

### Supplementary Fig2:

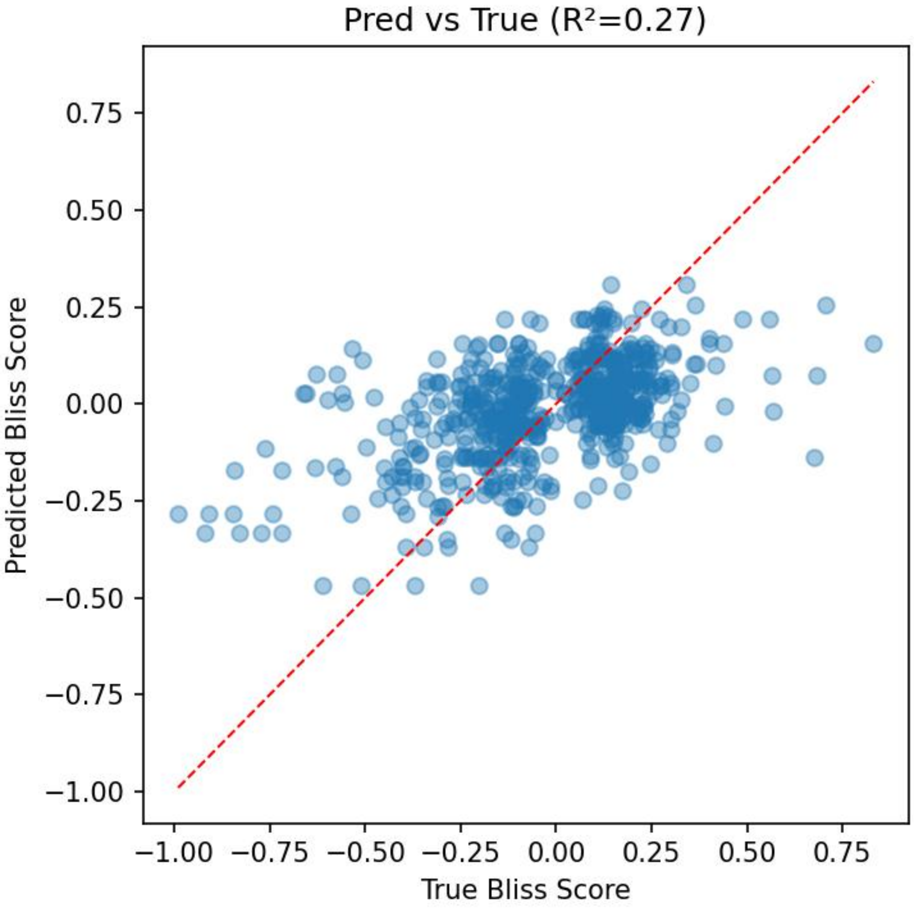


**Supplementary Figure 2.**

*Predicted vs. true Bliss ε values for the regression baseline (exp08). Each point represents a drug–pair–strain sample in the CV1 outer test set. Values cluster densely around the additive region (−0.1 to +0.1), and regression performance is strongly affected by noise in this range ($R^2$ = 0.27). This pattern motivated the use of a margin-based threshold ($|\varepsilon| \leq 0.05$) to exclude ambiguous additive interactions during binary-label construction.*
