## Supplementary Methods for "Bioactivity-Driven Prediction of Antibacterial Synergy Using Machine Learning Models"

**S1. Detailed description of feature representations**

We evaluated multiple strategies for constructing pairwise drug features from Chemical Checker (CC) embeddings to ensure that the final representation used in the main experiments was optimal.

1. **Concatenation (DrugA ⊕ DrugB)**
   Each drug's raw 25 × 128 CC signature was concatenated into a 6,400-dimensional vector. Although information-rich, this approach was order-sensitive and prone to overfitting in preliminary CV1 tests.

2. **Compact similarity vectors**
   Cosine similarity and Euclidean distance were computed at the coarse CC subspace level (25 domains, yielding 50 features). This largely discarded fine-grained structure and underperformed.

3. **Elementwise similarity (final choice)**
   For every CC dimension (25 × 128), we computed pairwise cosine similarity and Euclidean-derived similarity. The resulting 6,400 features preserved subspace-specific structure and consistently outperformed alternatives under CV1 and CV2.

**S2. Feature-selection pipeline (exp05 family)**

To reduce dimensionality without degrading performance:

- A LightGBM classifier was trained on the full 6,400-feature elementwise space.
- Gain importances were computed across all tree splits.
- Features exceeding a cumulative gain threshold were retained.
- Reduced sets were validated on representative CV1 and CV2 splits.

Before computing gain-based importances, we removed zero-variance features and discarded features whose absolute Pearson correlation with the binary interaction label on the CV1 training subset (grouped by drug pair) was < 0.01. This yielded a stable 1,578-feature subset used for all subsequent HALO-CV1 experiments.

**S3. Additional model variants**

**S3.1 Neural-network baseline (exp10 / exp10b)**

A feed-forward neural network comprising 2–3 dense layers with batch normalization, ReLU activation, and dropout was implemented. Training followed the same nested CV1 scheme. Both NN variants displayed overfitting and underperformed LightGBM, confirming that tree-based models were better suited for the dataset's structure.

#### S3.2 HALO-CV1s, HALO-CV1, and HALO-CV2s implementations

All variants shared identical LightGBM hyperparameter-search procedures. They differed only in grouping strategies:

- **HALO-CV1s:** test folds contain unseen drug pairs.
- **HALO-CV1:** same split as **HALO-CV1s** but trained without strain embeddings.
- **HALO-CV2s:** test folds contain unseen strains and unseen drug pairs; all cross-edge rows were removed to eliminate leakage.

#### S3.3 Lenient baseline: Standard stratified CV

For compatibility with earlier literature, we evaluated a conventional 5-fold stratified CV that does not enforce pair-level grouping. This method is known to inflate scores due to drug-identity leakage across folds.

#### S3.4 Multiclass and regression formulations

We tested two alternative problem formulations:

- **Multiclass classification:** (synergy, additive, antagonism) using the original Bliss categories. Performance was unstable due to poor separability of additive values.
- **Regression:** predicting continuous ε. Noise near the additive boundary reduced correlation with measured values, resulting in weaker generalization under both CV1 and CV2.

Given these limitations, all main analyses used the binary synergy–antagonism task.

### S4. Hyperparameter-search details

LightGBM models were tuned using nested 3-fold inner CV with early stopping. Randomized search covered learning rate, depth, number of leaves, feature and data subsampling, and regularization.
Neural-network baselines used a fixed small grid over hidden-layer size, dropout, and learning rate. Supplementary Table 1 reports the complete specification of every experiment, including all configurations and resulting evaluation metrics.

### S5. Additional procedural details

In the INDIGO compendium, quantitative Loewe α interaction scores were derived from 4×4 checkerboard assays measuring *E. coli* growth across a grid of drug concentrations for each pair. Deviations from Loewe additivity were converted to a continuous α score, where $\alpha < -0.5$ indicates synergy and $\alpha > 1$ indicates antagonism; these thresholds correspond approximately to Bliss ε scores of ≈ −0.3 (synergy) and +0.3 (antagonism). The INDIGO panel included 171 antibacterial pairs spanning aminoglycosides, quinolones, β-lactams, macrolides, and protein-synthesis inhibitors. Compounds were mapped to standardized InChIKeys, and non-antibacterial stress agents (e.g., $H_2O_2$) were excluded.

Chemical Checker signatures were retrieved using the official API. Random seeds were fixed across preprocessing, data-splitting, hyperparameter sampling, and model training for reproducibility. All scripts for dataset construction, feature engineering, and modeling will be released publicly upon publication.

**AI assistance details:** Large language models (ChatGPT 5.1 and Opus 4.5) were used for grammar correction and clarity editing. No scientific content, results, figures, or references were generated by AI.

**References:**


1. Species-specific activity of antibacterial drug combinations | Nature [Internet]. [cited 2025 Nov 30]. Available from: https://www.nature.com/articles/s41586-018-0278-9

2. Cacace E, Kim V, Varik V, Knopp M, Tietgen M, Brauer-Nikonow A, et al. Systematic analysis of drug combinations against Gram-positive bacteria. Nat Microbiol. 2023 Nov;8(11):2196–212.

3. Frontiers | ACDB: An Antibiotic Combination DataBase [Internet]. [cited 2025 Nov 30]. Available from: https://www.frontiersin.org/journals/pharmacology/articles/10.3389/fphar.2022.869983/full

4. Chandrasekaran S, Cokol-Cakmak M, Sahin N, Yilancioglu K, Kazan H, Collins JJ, et al. Chemogenomics and orthology-based design of antibiotic combination therapies. Mol Syst Biol. 2016 May 23;12(5):872.
